# Dissociation of CED-4 from CED-9 upon EGL-1 binding: Molecular mechanism of linear apoptotic pathway in *Caenorhabditis elegans*

**DOI:** 10.1101/2022.06.01.494320

**Authors:** C. Narendra Reddy, Ramasubbu Sankararamakrishnan

**Affiliations:** Department of Biological Sciences and Bioengineering, Indian Institute of Technology Kanpur, Kanpur 208016, Uttar Pradesh, India; Mehta Family Centre for Engineering in Medicine, Indian Institute of Technology Kanpur, Kanpur 208016, Uttar Pradesh, India

## Abstract

Many steps in programmed cell death are evolutionarily conserved across different species. The *Caenorhabditis elegans* proteins CED-9, CED-4 and EGL-1 involved in apoptosis are respectively homologous to anti-apoptotic Bcl-2 proteins, Apaf-1 and the “BH3-only” pro- apototic proteins in mammals. In the linear apoptotic pathway of *C.elegans*, EGL-1 binding to CED-9 leads to the release of CED-4 from CED-9/CED-4 complex. The molecular events leading to this process are not clearly elucidated. While the structures of CED-9 apo, CED- 9/EGL-1 and CED-9/CED-4 complexes are known, the CED-9/CED-4/EGL-1 ternary complex structure is not yet determined. In this work, we modeled this ternary complex and performed molecular dynamics simulations of six different systems involving CED-9. CED-9 displays differential dynamics depending upon whether it is bound to CED-4 and/or EGL-1. CED-4 exists as an asymmetric dimer (CED4a and CED4b) in CED-9/CED-4 complex. CED-4a exhibits higher conformational flexibility when simulated without CED-4b. Principal Component Analysis revealed that the direction of CED-4a’s winged-helix domain motion differs in the ternary complex. Upon EGL-1 binding, majority of non-covalent interactions involving CARD domain in the CED-4a-CED-9 interface have weakened and only half of the contacts found in the crystal structure between α/β domain of CED4a and CED-9 are found to be stable. Additional stable contacts in the ternary complex and differential dynamics indicate that winged-helix domain may play a role in CED-4a’s dissociation from CED-9. This study has provided a molecular level understanding of potential intermediate states that are likely to occur at the time of CED-4a’s release from CED-9.

## Introduction

Programmed cell death (PCD) is a physiologically important process and crucial in maintaining the homeostasis during different stages of development ^1, 2^. Several steps in PCD are conserved across diverse organisms such as *Caenorhabditis elegans*, *Drosophila melanogaster* and humans ^3^. There are also unique features of PCD in species groups like nematodes ^4^. Biochemical and genetic studies revealed the role of four genes in regulating the cell death in *C. elegans* ^5–7^. The corresponding proteins of these genes are CED-9, CED-4, EGL-1 and CED-3. CED-9 in *C. elegans* is homologous to mammalian anti-apototic Bcl-2 proteins and the experimentally determined structure showed that CED-9 adopts the characteristic Bcl-2 helical fold found in mammalian counterparts ^8–10^. Experimental and computational studies have used the typical Bcl-2 helical fold of anti-apoptotic Bcl-2 proteins to target and design inhibitors that can antagonize the function of these proteins ^11–17^. Such molecules, called BH3-mimetic, are potential anti-cancer drugs and many of them have entered clinical or pre-clinical trails ^18–20^. CED-4, a homolog of human apoptotic activating factor 1 (Apaf1), is a pro-apoptotic adaptor protein and is sequestered by CED-9 bound to mitochondria ^21^. This prevents CED-4 to activate the inactive zymogen caspase CED-3. EGL-1 is a “BH3-only” pro-apoptotic protein which helps to release CED-4 bound to CED-9 and this process eventually activates the cysteine protease CED-3 ^5, 22, 23^. Structural and biochemical studies reveal the region important for recognition of CED-3 by CED-4 apoptosome and the functional importance of this interface region ^24^. The structure of CED-9 in complex with CED-4 has also been determined ^25^. CED-4 with nearly 550 residues has four domains, namely CARD, an α/β fold domain resembling P-loop NTPases, a helical domain and a winged-helix domain. The CED-9/CED-4 complex structures revealed several interesting features including the details of interacting interface. Although buried interface area in the complex is significant, poor shape complementarity and lack of many intermolecular interactions by the buried amino acid residues raises many interesting questions about the mode of CED-4’s dissociation from CED-9 after EGL-1 binding. Structure of CED-9 in complex with EGL-1 has been determined and showed a strong binding between the two molecules ^9^. The CED-9 in apo form and in complex with CED-4 are structurally very similar ^8, 25^ while the CED-9 in complex with EGL-1 exhibits noticeable differences from the other two forms. Upon binding, EGL-1 induces specific conformational changes in CED-9 which destabilizes the CED-9/CED-4 complex. Using yeast-based systems, it has been shown that human Bcl-2 binds to EGL-1 which prevents its binding to CED-9 and suppress the cell death in *C. elegans* ^26^. Molecular dynamics (MD) simulation studies of CED-9 in apo form revealed that the helices forming hydrophobic pocket is rigid compared to the mammalian anti-apoptotic Bcl-2 proteins ^27^.

The structural studies combined with mutagenesis and biochemical approach have elucidated some important conclusions and these are vital for understanding the linear pathway from EGL-1 to CED-3 that leads to apoptosis. Some of the important points from earlier studies include two distinct binding regions for CED-4 and EGL-1 in CED-9 ^9, 25^. The binding affinity of EGL-1 for CED-9 is 8 fold higher than that of CED-4 ^25^. Comparison of CED-9 structures in apo and EGL-1-bound forms indicate the movement of a specific α-helix in CED-9 towards the direction of CED-4 by nearly 6 Å ^25^. This conformational change seems to be important for the dissociation of CED-4 from CED-9. Some important residues involved in CED-9/EGL-1 interactions and interactions between CED-9 and CED-4 have been identified ^25^. To understand the activation of inactive zymogen CED-3 by CED-4, the molecular events that lead to the disruption of CED-4 interactions with CED-9 upon EGL-1 binding have to be clearly understood. Although the structures of CED-9 in apo-form, EGL-1-bound and CED-4-bound forms are available, the ternary structure in which EGL-1 and CED-4 simultaneously bound to CED-9 is not yet determined. It is possible that such ternary complex cannot be captured as EGL-1 binding CED-9 is likely to result in the immediate release of CED-4. To investigate these points, we have performed molecular dynamics (MD) simulations of free CED-9 and in complex with EGL-1 and CED-4. We have also built a ternary complex of EGL-1/CED-9/CED-4 and carried out MD simulations to investigate the stability and eventual release of CED-4. We have analyzed CED-9’s dynamic behavior in all four forms in 500 ns to more than one μs simulations and it clearly showed that the EGL-1 and CED-4 proteins have different influences on CED-9 after the complex formation. Interactions between CED-4 and CED-9 have significantly weakened in the ternary complex in which EGL-1 is also bound to CED-9. Our studies indicate the possible involvement of winged helix domain of CED-4 although it does not directly interact with CED-9. Our results have furthered our knowledge in understanding the cell death pathway in *C. elegans* which will have implications in understanding the molecular mechanism of apoptosis in other organisms such as *D. melanogaster* and humans.

## Materials and Methods

### Experimental structures selected for the study

The experimentally determined structures of CED-9 in apo form (PDB ID: 1OHU) ^8^, in complex with EGL-1 (PDB ID: 1TY4) ^9^ and in complex with CED-4 (PDB ID: 2A5Y) ^25^ were downloaded from the Protein Data Bank ^28^. Selenomethionine residues in the crystal structures of CED-9 apo and CED-9/EGL-1 complex structures were substituted back to methionine using Chimera ^29^. The unit cell of CED-9/EGL-1 (PDB ID: 1TY4) has two complex structures. We selected Chain A and Chain C as starting structures for molecular dynamics simulations of CED- 9/EGL-1 complex. Pro-148 in Chain A of CED-9/EGL-1 was substituted to the wild-type residue Leu using Chimera. Similarly, serine residues at positions 107, 135 and 164 were reverted back to Cys in CED-9/CED-4 complex structure. The missing regions of CED-9/CED-4 complex (Chain A: 161-162; Chain B: 311- 312, 417-423, 488-520; Chain C: 415-426, 483-532) were modeled using loop class of Modeller 9.14 software ^30–32^. In the case of CED-9 in complex with CED-4, CED-4 is present as an asymmetric dimer. Hence, we considered two different systems, CED-9 with CED-4 dimers (CED-4a and CED-4b) and also CED-9 with just CED-4a. These are respectively designated as CED-9/CED-4 and CED-9/CED4a. It should be noted that in the case of CED-9 with CED-4 dimers, only CED-4a is involved in interactions with CED-9 as shown by the crystal structure ^25^. As far as the CED-9 in complex with the monomer CED-4a, we simply removed the other monomer CED-4b which is involved in contact with only CED-4a and this was done to see the effect of CED-4 dimers in CED-9 complex stability.

### Modeling of EGL-1/CED-9/CED-4 complex

The ternary complex structure of CED-9/CED-4/EGL-1 has not yet been determined experimentally and it might be due to the transient nature of the complex. We used the experimentally determined complex structures CED-9/EGL-1 and CED-9/CED-4 to build this ternary complex. These structures were used as template structures to model using the software Modeller 9.14 ^30–32^. With CED-4 and EGL-1 missing respectively in CED-9/EGL-1 and CED- 9/CED-4 complex structures, the alignment of the target and template sequences were done as follows. The target sequence (CED-9/CED-4/EGL-1 complex) and the sequences of the template structures (CED-9/CED-4 and CED-9/EGL-1) were aligned by stacking the identical regions of amino acid sequences and gaps were introduced in the missing regions of the corresponding structures. This target-template alignment was used to produce five best models and the model with the lowest DOPE (Discrete Optimized Protein Energy) score ^33^ was selected for modeling the loops. Ten models with different loop conformations were generated and again the model with the lowest DOPE score was selected for further studies. The generated CED-9/CED-4/EGL- 1 ternary complex is shown in Figure 1. As in the case of CED-9/CED-4 complex, we considered the ternary complex in two different forms, one with CED-4b and the other without CED-4b, referred respectively as CED-9/CED-4/EGL-1 and the CED-9/CED-4a/EGL-1 ternary complexes. The coordinates of CED-9/CED-4/EGL-1 ternary complex are provided in the Supplementary Material.

**Figure 1:**
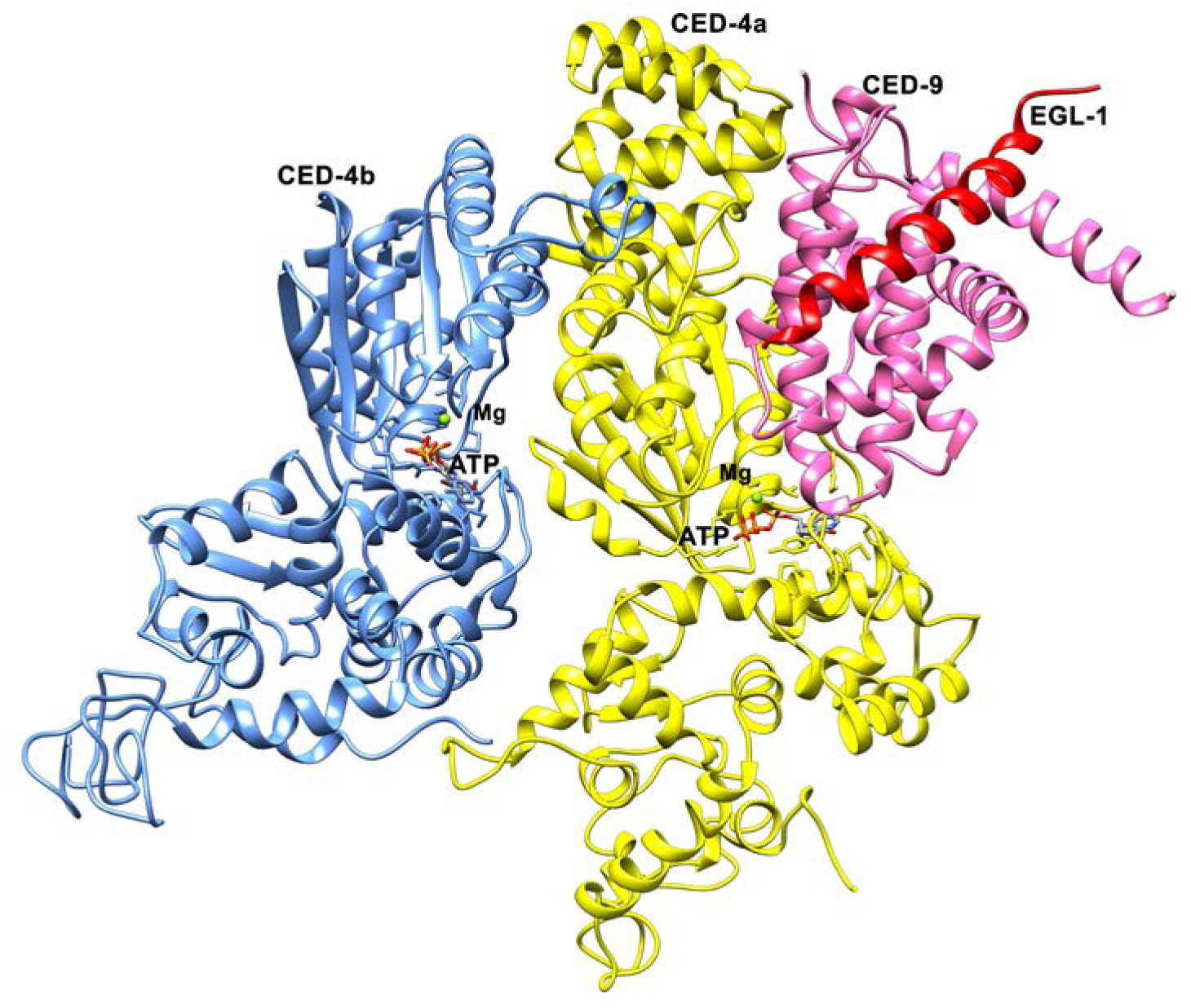
Model of CED-4/CED-9/EGL-1 ternary complex generated from the experimentally determined complex structures of CED-9/CED-4 (PDB ID: 2A5Y) and CED-9/EGL-1 (PDB ID: 1TY4). EGL-1, CED-9, CED-4a and CED-4b are represented in red, pink, yellow and blue colors respectively. The bound ATP and Mg2+ ions are displayed in the structure in stick and sphere representations respectively. The coordinates of the model are available in the Supplementary Information.

### Molecular dynamics simulations of CED-9 and its complex structures

All simulations were carried out using Gromacs 5.1.4 version ^34, 35^ with CHARMM36 force-field ^36, 37^. For each system simulated, the following protocol was used. The starting structure was placed inside a cubic box such that the minimum distance between the protein and the box was at least 10 Å. The structure was then solvated with TIP3P water ^38^ and the entire system was made neutral by adding required number of Na+ and Cl- ions. For both van der Waals and columbic interactions, the cutoff distance used for evaluating the short-range interactions was 10 Å. Particle Mesh Ewald (PME) method ^39^ was applied to calculate long-range electrostatic interactions. The entire system was energy minimized using steepest descent method till the maximum force on any particle was less than 500 kJ/mol/nm. The system was then brought to room temperature (300 K) by carrying out an NVT simulation for about 1 ns. During the NVT simulation, the system was coupled to the V-rescale thermostat ^40^ with the coupling constant of 0.1 ps. After properly equilibrating the system in NVT ensemble, it was further equilibrated using NPT ensemble. The system was coupled with the Parrinello-Rahman barostat ^41^ at a coupling constant of 0.2 ps with a reference pressure 1 atm. After performing further equilibration for a period of 1 ns at 300 K, the pressure in the system settled around 1 atm. Protein was restrained with the harmonic potential of 1000 kJ/mol during both stages of equilibrations. It must be noted that few of the modeled missing loops of CED-4 in both CED- 9/CED-4 and CED-9/CED-4/EGL-1 complex structures are very long. To sample enough conformations and to make sure that the loops adopt the best conformations with respect to the rest of the protein, we ran 25 ns restrained simulation of both the complexes by restraining the rest of the protein. This is followed by a production run for a period of 500 ns to more than 1μs with no restraints. Similar steps were taken while simulating CED-9/CED-4a and CED-9/CED- 4a/EGL-1 systems. In the case of CED-9 apo and CED-9/EGL-1, the trajectories generated were 500 ns in length. The production runs for CED-9/CED-4 and CED-9/CED-4/EGL-1 complexes were of one or more than one µs in length. Table 1 provides the details of all the simulations performed in this study.

**Table 1:** Summary of the simulations carried out in this study

| System | Description | Starting structure | No. of atoms | Production run (in ns) |
| --- | --- | --- | --- | --- |
| System-1 | CED-9 apo | PDB ID: 1OHU | 35629 | 500 |
| System-2 | CED-9/EGL-1 | PDB ID: 1TY4 | 54480 | 500 |
| System-3 | CED-9/CED-4a | PDB ID: 2A5Y-<br>ΔCED-4b<sup>a</sup> | 217599 | 500 |
| System-4 | CED-9/CED-4a/EGL-1 | Model<sup>b</sup>-ΔCED-4b<sup>a</sup> | 219525 | 500 |
| System-5 | CED-9/CED-4 | PDB ID: 2A5Y | 327643 | 1000 |
| System-6 | CED-9/CED-4/EGL-1 | Model<sup>b</sup> | 351975 | 1080 |

<sup>a</sup>CED-4b structure was removed from the experimentally determined structure
<sup>b</sup>The ternary complex was modeled. For details of modeling, see the Method section
<sup>c</sup>CED-4b structure was removed from the model

### Principal Component Analysis (PCA)

In PCA, the first few principal modes of covariance matrix computed from the trajectory contribute 70 to 90% of the fluctuations of the molecule. The space spanned by high eigenvalue principal modes is called essential space, and it is believed to contain functional collective motions. The covariance matrix and the eigenvectors of the matrix were calculated after utilizing the Gromacs covar utility. The analysis of eigenvectors and the projection of the trajectory onto the eigenvectors were done using anaeig utility of Gromacs. The essential space of the system of interest was derived by projecting the discrete trajectories of the concatenated trajectory onto the first two eigenvectors. For example, CED-9 concatenated trajectory was obtained by concatenating the last 500 ns trajectories of System-1, System-2, System-5 and System-6.

## Results

### Dynamics of CED-9 in different simulated systems

We have carried out MD simulations of six different systems involving CED-9, CED-4 and EGL-1 and the summary of simulations is provided in Table 1. Different properties of the simulated systems were analyzed. First we wanted to see how CED-9 dynamics is affected when it is bound to EGL-1 and/or CED-4. Association of CED-9 with EGL-1, CED-4 and with both of them might have different kinds of influence in inducing conformational changes in CED-9. We calculated the root mean square deviation (RMSD) of CED-9 with reference structures corresponding to the initial structures of different systems and only Cα atoms were considered for this purpose (Table 2).

**Table 2:**
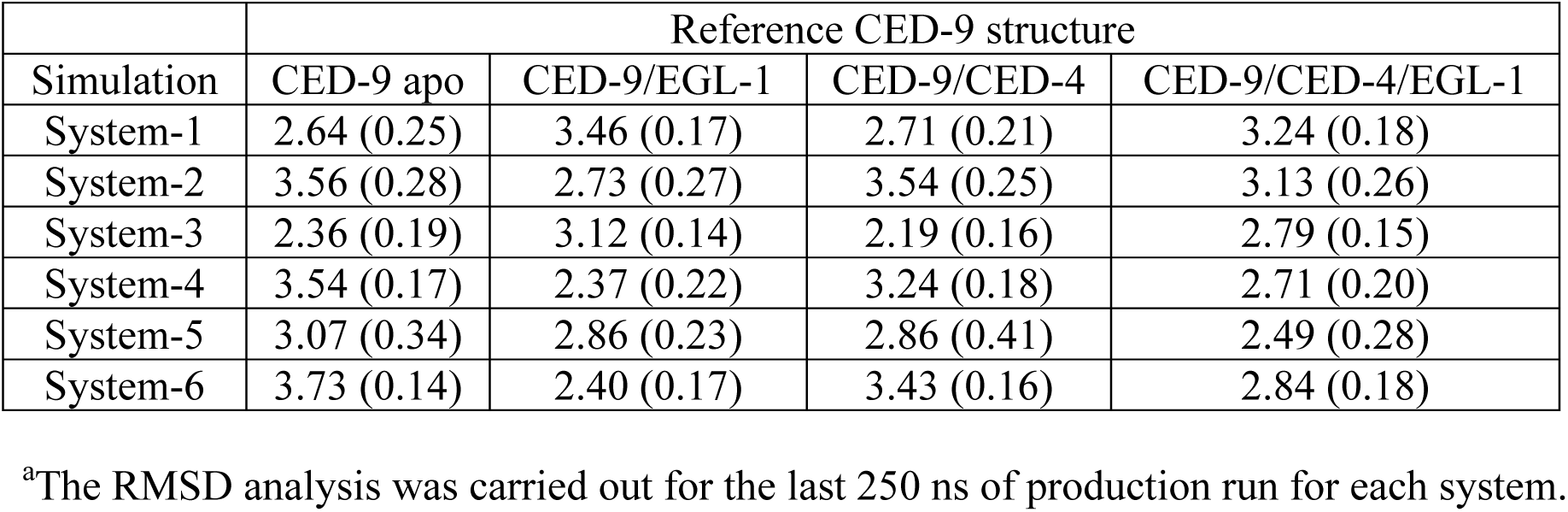
Average Root Mean Square Deviation (standard deviation) (in Å)^a^ of CED-9 in different simulated systems

When the reference structure is CED-9 apo, it exhibited highest average RMSD (denoted as <RMSD>) with the simulated structures of System-2 (CED-9 in complex with EGL-1; <RMSD> = 3.56 Å) or System-6 (CED-9 which is part of the ternary complex; <RMSD> = 3.73 Å). This can be understood as the crystal structures of CED-9 in apo state and CED-9 in complex with EGL-1 show significant differences especially in the hydrophobic binding pocket ^9^. A similar observation was made if we consider CED-9 when it is in complex with EGL-1 as the reference structure. The maximum deviation was observed when CED-9 of CED-9/EGL-1 was compared with the simulated structures of System-1 (CED-9 apo; <RMSD> = 3.46 Å). When CED-9 in complex with CED-4 asymmetric dimer was considered as reference structure, as anticipated, the maximum deviation was observed with the simulated structures of System-2 (CED-9 in CED-9/EGL-1 complex; <RMSD> = 3.54 Å). However, it is also interesting to note that CED-9 in CED-9/CED-4 complex also exhibits higher deviation in System-6 simulation as CED-9 was part of the ternary CED-9/CED-4/EGL-1 complex (<RMSD> = 3.43 Å). Differences in the RMSD analysis are observed when simulations of System-3 (CED-9/CED-4a) and System-5 (CED-9/CED-4) were compared indicating that CED-4 whether it is present as an asymmetric dimer or monomer in the CED-9/CED-4 complex seems to exert differential effects on CED-9 structure. This is in spite of the fact that CED-4b in the asymmetric dimer is not directly in contact with CED-9. When CED-9 as part of the ternary complex (CED-9/CED- 4/EGL-1 or CED-9/CED-4a/EGL-1) was considered as the reference structure, it showed higher structural deviations with simulated structures of both System-1 (CED-9 apo; <RMSD> = 3.24 Å) and System-2 (CED-9/EGL-1 complex; <RMSD> = 3.13 Å) indicating that CED-9 in ternary complex has features of both CED-9 apo and CED-9/EGL-1 complex.

RMSD indicates the overall deviation of MD simulated structure from the reference structure and does not reveal which residue or region exhibits more flexibility. Hence, we also plotted root mean-square fluctuation (RMSF) which gives a measure of average deviation of each residue from its reference position over a period of time. RMSF plot of CED-9 simulations with respect to CED-9 apo structure is shown in Figure 2. Residue range of helical segments in CED-9 as defined in the crystal structure is given in Table S1. RMSF plot with CED-9 as initial structure from CED-9/EGL-1 exhibits similar behavior (Supplementary Figure S1). It is clear from the plot that loop regions are generally flexible. Two particular loops show very high fluctuations in at least one of the simulations. The loop connecting the first two helices is long (residues 95-109; residue numbering according to PDB ID: 1OHU) and is known to be flexible. However, the loop connecting the α4 and α5 helices also display large flexibility and it is especially pronounced in the simulations of System-2 (CED-9/EGL-1) and System-5 (CED- 9/CED-4). It is interesting to note that these loops form part of the interface between CED-9 and CED-4 in the complex structure. Superposition of MD simulated structures saved at the end of production runs from different simulations on the CED-9 apo crystal structure is shown in Figure 3 and the two loops undergoing large conformational changes are highlighted in the figure.

**Figure 2:**
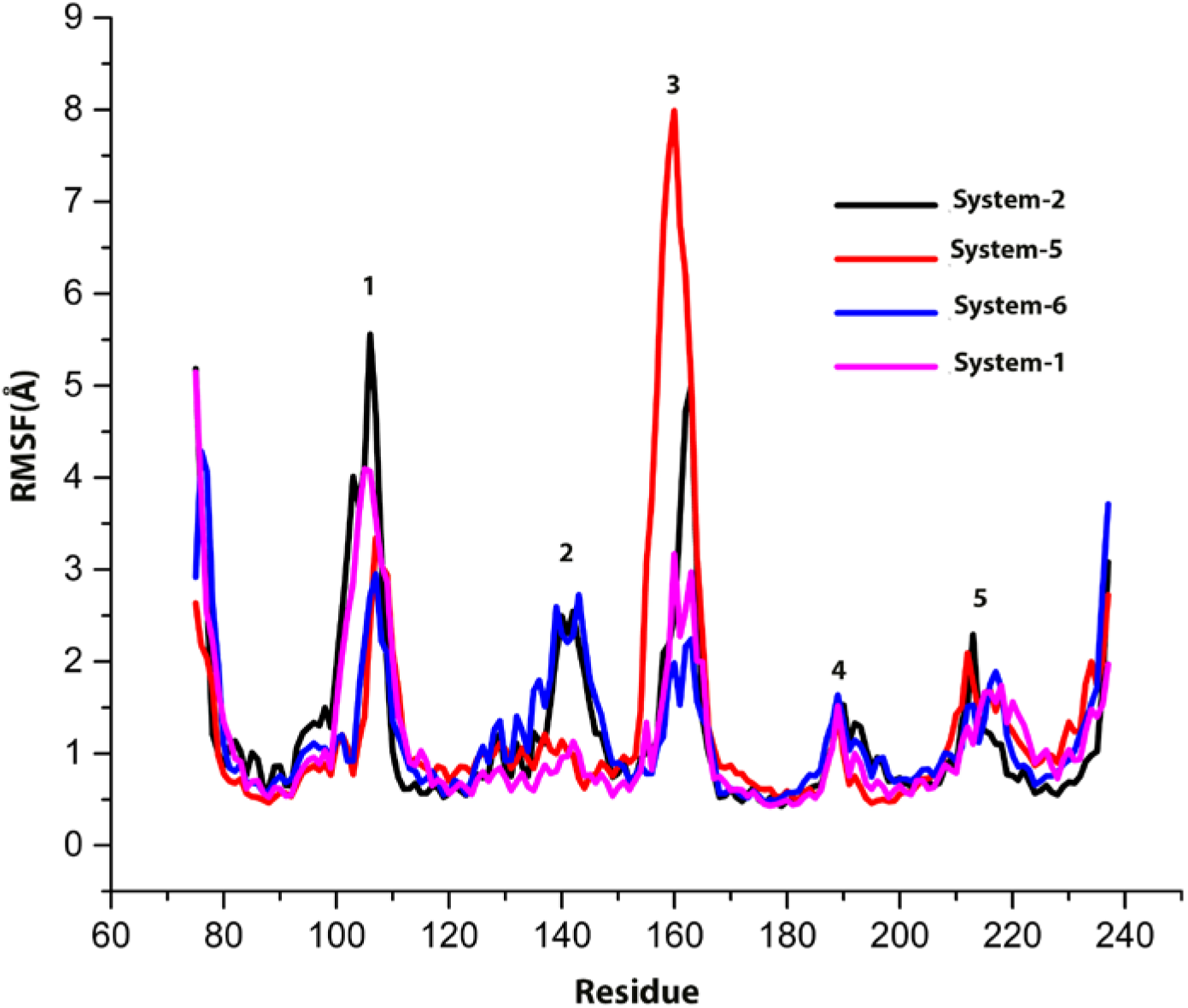
Root mean square fluctuation (RMSF) of CED-9 in different simulated systems calculated with CED-9 Apo as the reference structure. The CED-9 regions that undergo large fluctuations are the loops connecting helices (1) α1 and α2 (95-109), (2) α3 and α4 (residues 130-150), (3) α4 and α5 (residues 155-166), (4) α5 and α6 (residues 185-194) and (5) α6 and α7 (residues 205-220).

**Figure 3:**
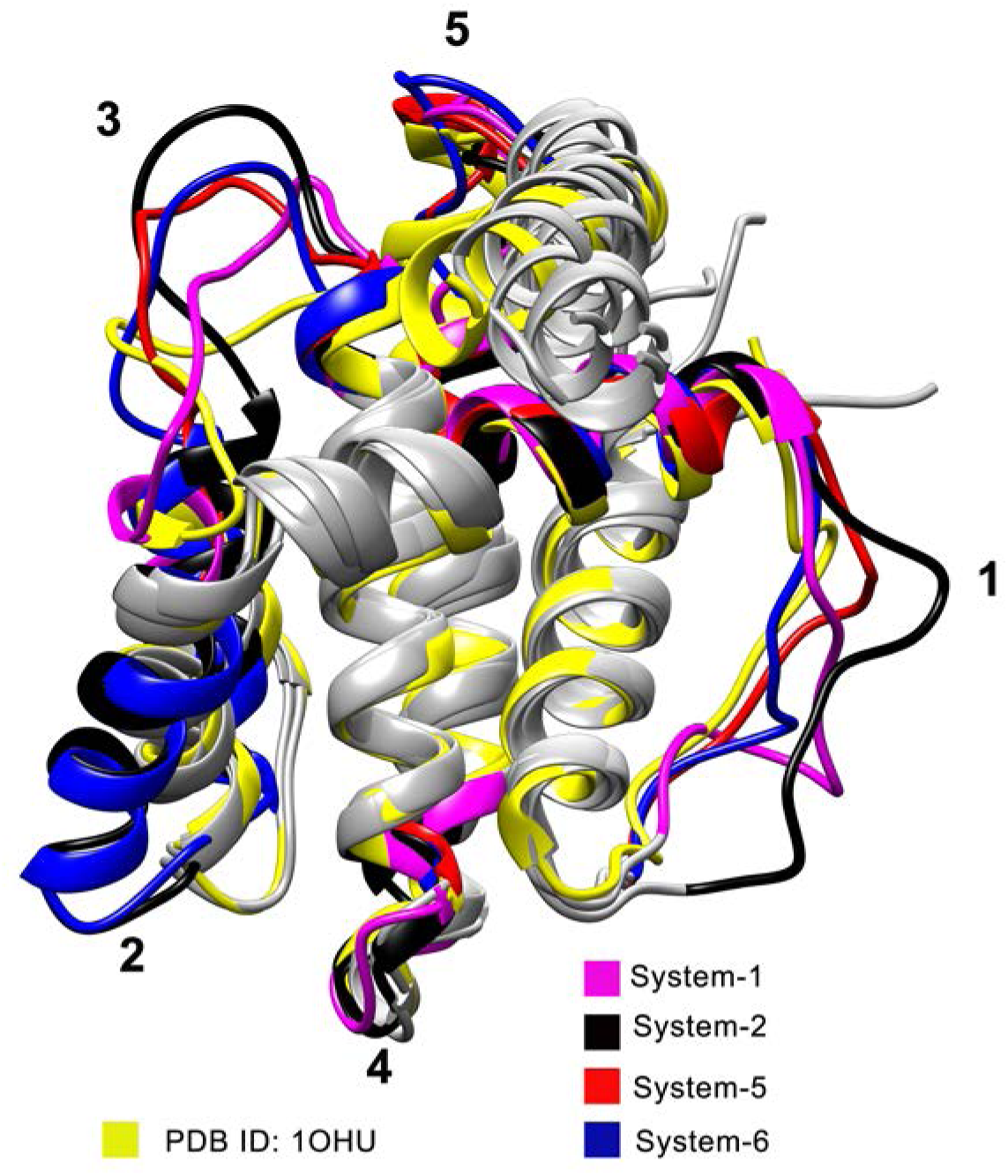
MD simulated structures of CED-9 saved at the end of production runs for each system are superposed on the crystal structure (PDB ID: 1OHU). The regions with large conformational fluctuations are labeled and the residues corresponding to these regions are given in Figure 2 caption.

### 2D-RMSD Plots of CED-9

As CED-9 is simulated in different conditions, we also analyzed the 2D-RMSD plots for various simulations. 2D-RMSD plots differ from the usual RMSD analysis in the sense that in 2D-RMSD analysis, each MD-simulated structure is compared with every MD snapshot of a particular simulation. This will give us an idea of how much the conformational space of CED-9 is sampled in a given simulation under a particular condition. Comparison of 2D-RMSD plots of different CED-9 simulations will reveal in which simulation CED-9 sampled larger conformational space. We have used only Cα atoms of CED-9 for this purpose. The 2D-RMSD plots of CED-9 are shown for System-1 (CED-9 apo), System-2 (CED-9/EGL-1), System-5 (CED-9/CED-4) and System-6 (CED-9/CED-4/EGL-1) simulations (Figure 4). The 2D-RMSD plots of other two systems (System-4 and System-5) are provided in the Supplementary Information (Figure S2). Among the six simulations, CED-9 apo exhibits the least conformational fluctuations which was also reported in an earlier study from our lab ^27^. The 2D- RMSD plots of System-2, System-5 and System-6 show that the MD simulated structures at the beginning of the production runs deviate from those simulated at the end of the production runs. This is especially evident when CED-9 is associated with CED-4 as can be seen for System-5 (CED-9/CED-4) and System-6 (CED-9/CED-4/EGL-1) simulations. Maximum deviations were seen for CED-9 structures in System-5 (CED-9/CED-4) simulations around 800 ns as shown by the red color in the plot. When CED-9 is simulated with CED-4a without including CED-4b, both System-3 and System-4 indicate that structural deviations are found from the beginning within 100 ns and this is especially prominent in System-4 in EGL-1 bound form (CED-9/CED- 4a/EGL-1) again reiterating that the absence of CED-4b results in higher conformational fluctuations. In summary, the 2D-RMSD plots clearly demonstrate that the CED-9 undergoes larger conformational changes when it is in complex with CED-4.

**Figure 4:**
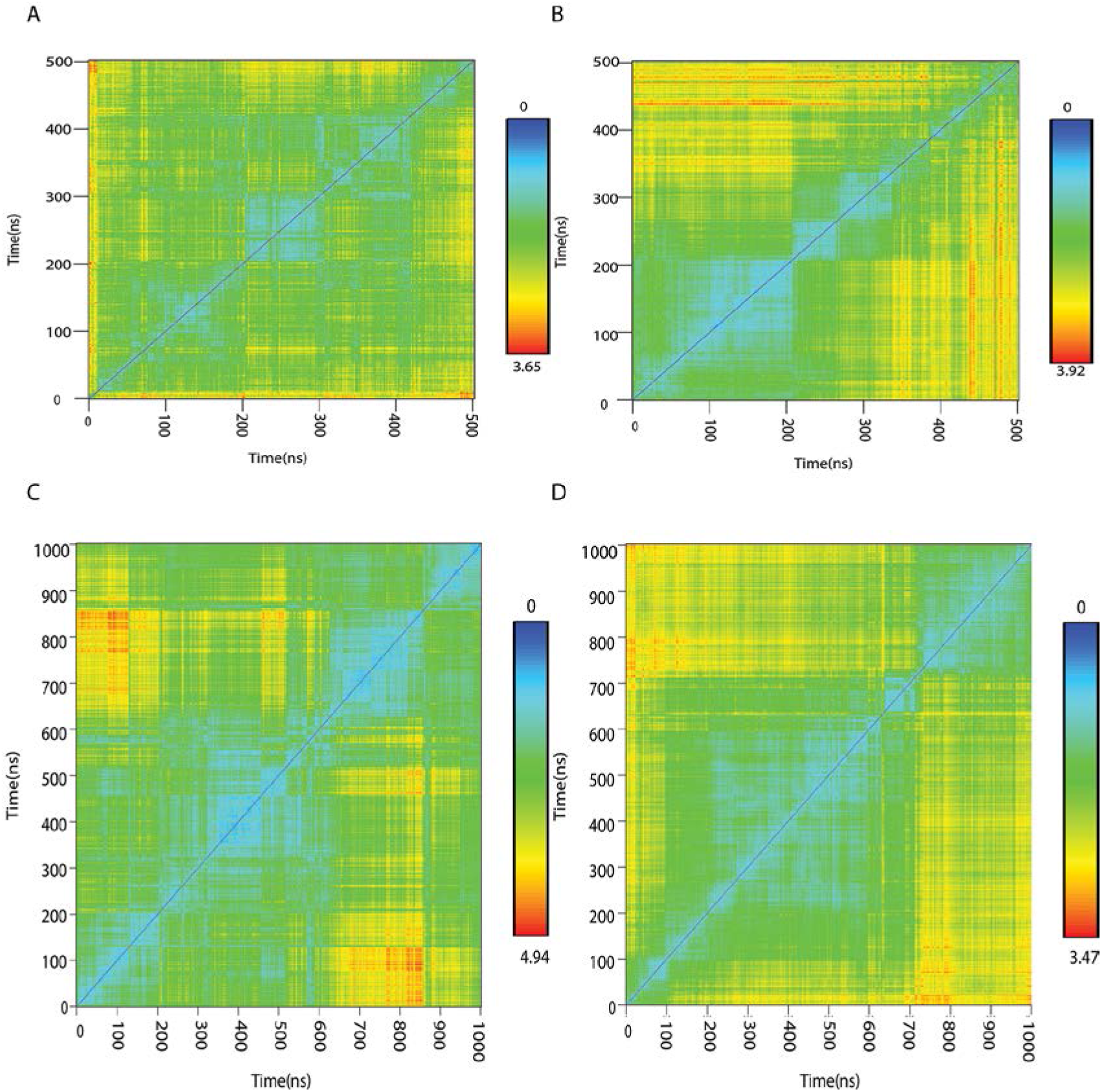
2D-RMSD plots of CED-9 in different simulations as a function of time shown for (A) System-I (Ced-9 apo), (B) System-2 (CED-9/EGL-1), (C) System-5 (CED-9/CED-4) and System-6 (CED-9/CED-4/EGL-1). RMSD values (in Å) are displayed from blue to red covering the range of lowest to highest values.

### Dynamics of CED-4 in the simulated systems

In the current study, CED-4 is considered as part of CED-9/CED-4 complex or CED- 9/CED-4/EGL-1 ternary complex. CED-4 is present as an asymmetric dimer (CED-4a and CED- 4b) when it is in complex with CED-9 in the experimentally determined structure ^25^. Only CED- 4a is in direct contact with CED-9. Although CED-4b is not interacting directly with CED-9, its interaction with CED-4a might have an indirect influence on CED-4a’s association with CED-9. Hence, we simulated both CED-9/CED-4 and CED-9/CED-4/EGL-1 complex structures with and without CED-4b (Table 1). We calculated RMSD of CED-4a (the monomer in association with CED-9) in different systems. The CED-4a in CED-9/CED-4 complex was considered as the reference structure and compared with the simulated structures of CED-9/CED-4 complex (Figure 5). When this complex was simulated without CED-4b in System-3 (CED-9/CED-4a), the average RMSD is much higher than that found in the simulations of System-5 (CED-9/CED- 4) in which CED-4b is present as asymmetric dimer. This immediately indicates that CED-4b which is interacting with CED-4a in CED-9/CED-4 complex is restricting its dynamics. A similar observation was made when the simulated structures of ternary complex were considered. In the simulations of System-6 (CED-9/CED-4/EGL-1), when CED-4b is present, the average RMSD of CED-4a is more than 3.5 Å lower than that found in the simulation of System-4 in which CED-4b was not included. This clearly indicates that CED-4b through its interactions with CED-4a reduces the conformational flexibility of CED-4a. Since CED-4 is present as an asymmetric dimer in its interaction with CED-9, we extended the simulations of only System-5 (CED-9/CED-4) and System-6 (CED-9/CED-4/EGL-1) up to 1 μs for further analysis. The RMSD trajectory remains stable for both the systems throughout the production runs (Figure 5).

**Figure 5:**
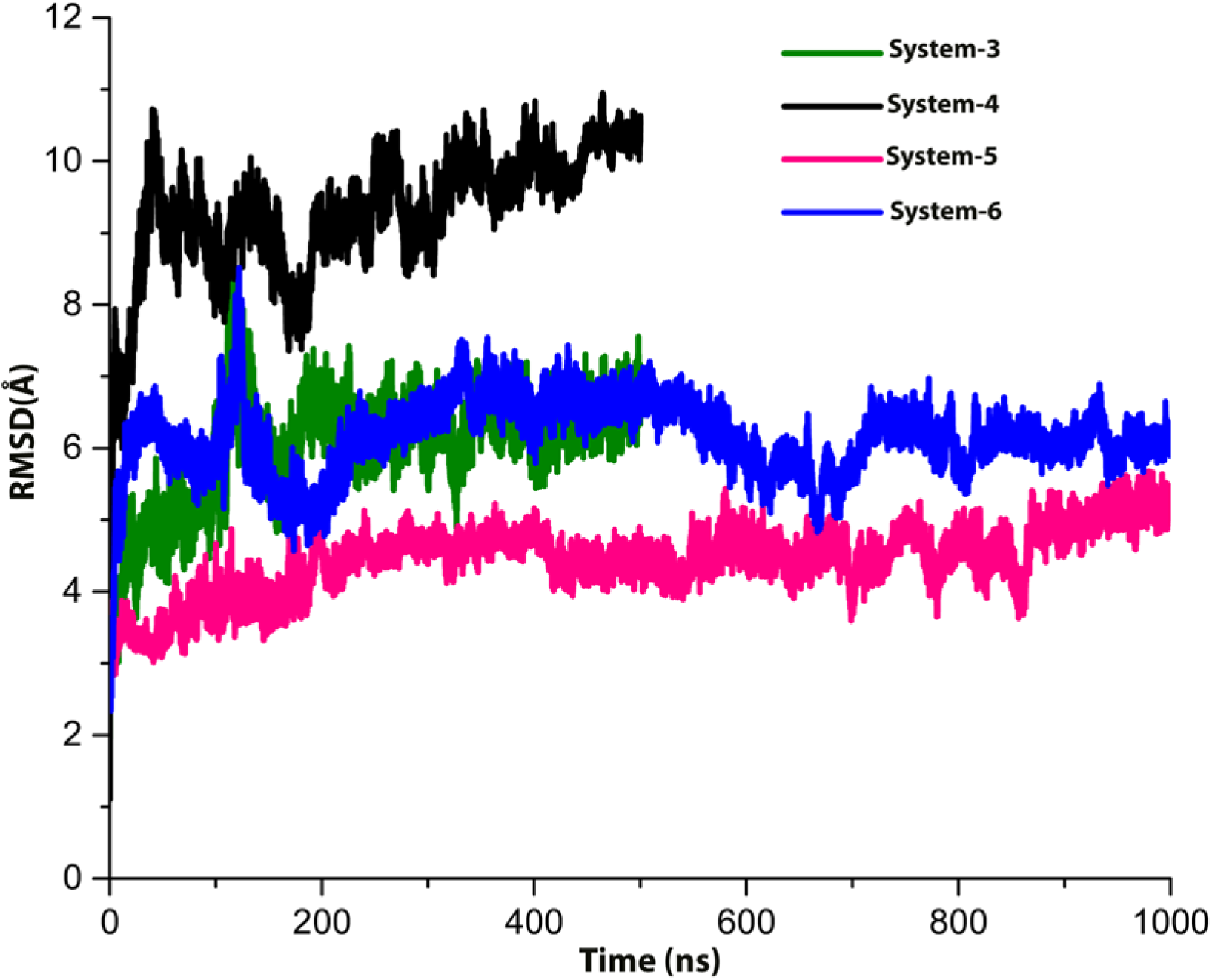
RMSD trajectories of CED-4a as a function of time in different simulations.

The RMSF plots of CED-4a in different simulations clearly indicate that the loops exhibit large fluctuations (Figure 6A). Different domains of CED-4 as defined in the crystal structure are given in Table S2. The long loop near the C-terminus (residues 479-535) especially shows conformational heterogeneity in all the simulations. Apart from the loops, we also see conformational transitions of a helix just before this loop (residues 470-478). In the simulation of System-6 in which the ternary complex with asymmetric CED-4 dimer was simulated, this helix seems to unwind with some loss of its helical character. The simulated structures saved at the end of the production runs from different simulations were superposed on the experimentally determined CED-4a monomer and the superposed structures are shown in Figure 6B. The regions showing large conformational plasticity are highlighted in the figure. As shown in the figure, the major conformational changes happen in the C-terminal winged helix domain (residues 371-549) of CED-4a. It is also the site of secondary binding site with CED-4b with which the residues from winged domain of CED-4a are involved in several hydrogen bond and salt-bridge interactions with the helical domain (residues 291-370) of CED-4b ^25^. This explains why the absence of CED-4b results in greater conformational changes of the winged helix domain of CED-4a (Figure 6). We also observe that the loop (residues 213-226) in the three-layered α/β fold in CED-4 undergoes fluctuations as evident from the RMSF plot. This loop is involved in interactions with the loop (residues 143-147) connecting third and fourth helices of CED-9 structure (see below).

**Figure 6:**
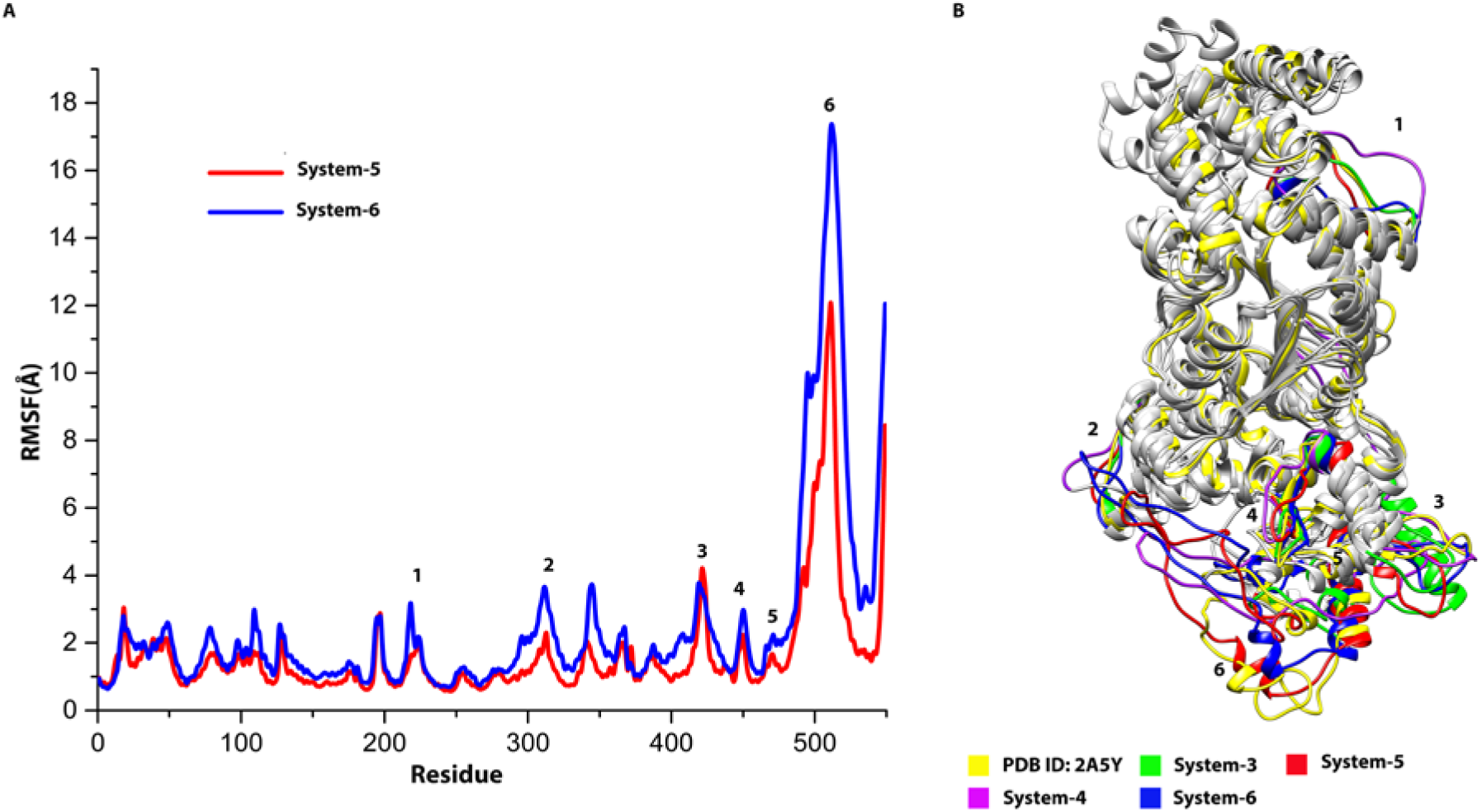
(A) RMSF analysis of CED-4a in System-5 (CED-9/CED-4) and System-6 (CED- 9/CED-4/EGL-1) simulations. RMSF fluctuations are higher for the ternary complex compared to System-5 simulation. (B) Superposition of MD simulated structures of CED-4a saved at the end of the production runs on the experimentally determined CED-4a structure (PDB ID: 2A5Y).

### Principal Component Analysis of CED-9 and CED-4 in different systems

Principal Component Analysis (PCA) has been used to understand the conformational dynamics of diverse proteins ^42–45^. We performed PCA to understand the atomic displacement and conformational flexibility of CED-9 and CED-4 in different conditions. For this purpose, we extracted different modes by performing PCA during MD simulations of systems focusing on the dynamics of either CED-9 or CED-4. The essential dynamics of the systems were retrieved as described in the Methods section and compared. The first two principal components (PC1 and PC2) dominate the overall motion and hence we have plotted these modes for CED-9 in different simulations (Figure 7). The plot PC1 vs PC2 clearly shows that CED-9 conformational changes are distinctly different in different conditions. Compared to the other three systems, CED-9 apo samples restricted conformational space. This was also demonstrated in earlier MD simulation studies in which the dynamics of CED-9 apo was compared with that of three mammalian anti-apoptotic homologs ^27^. CED-9 in complex with EGL-1 samples relatively larger conformational space with some overlap with CED-9 apo in the direction of PC2. However, CED-9/EGL-1 accesses different conformations in PC1 mode. The dynamics of CED-9 in complex with CED-4 and CED-9 as part of the ternary CED-9/CED-4/EGL-1 complex are distinct from CED-9 apo and CED-9/EGL-1 complex. The conformational space of CED-9 in CED-9/CED-4 and CED- 9/CED-4/EGL-1 displays some overlap along PC1 and exhibit distinct differences in the direction of PC2. Overall our PCA analysis clearly exhibit distinct conformational changes for CED-9 depending upon its interaction with CED-4 and/or EGL-1.

**Figure 7:**
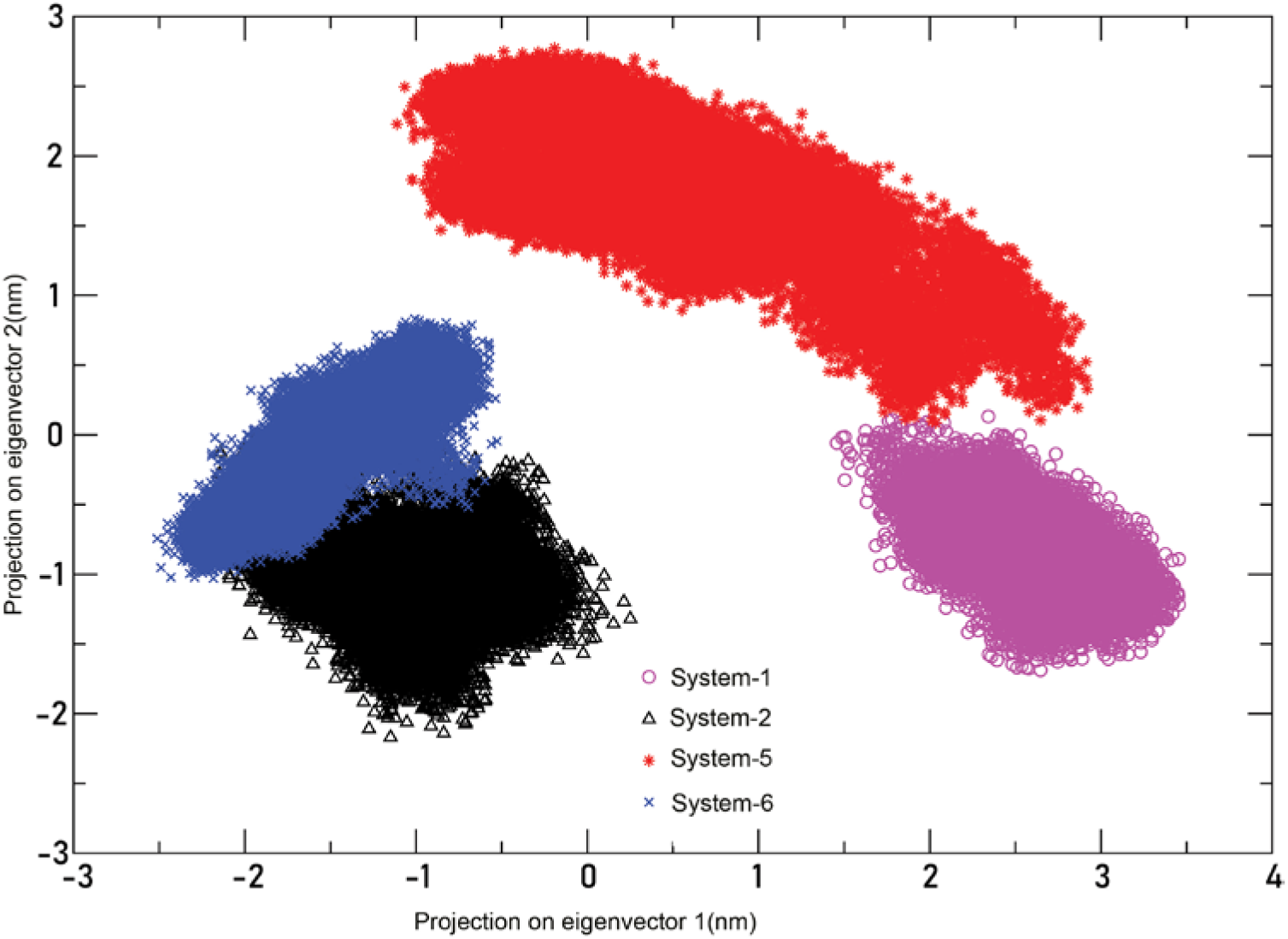
PC1 vs PC2 plot of CED-9 in different conditions. This analysis was done for the last 300 ns of production runs in each simulation

A similar PCA analysis was also performed on simulations involving CED-4a to understand the impact of binding to CED-9 and CED-9/EGL-1 (Figure 8). The PC1 vs PC2 plot shows that CED-4a undergoes distinct conformational changes when it is in complex with CED- 9 compared to its dynamics when CED-4a is part of the ternary CED-9/CED-4/EGL-1 complex. This is evident in the directions of both PC1 and PC2. To understand the conformational dynamics of CED-9/CED-4 complex with and without bound EGL-1, we plotted the porcupine plots of CED-9/CED-4 complex that describes the first eigenvector of PCA (Figure 9). The length and direction of arrows represent the motion density and motion direction of different domains/regions of the complex respectively. Porcupine plots of first mode of PCA of CED- 9/CED-4 complex were compared for the System-5 and System-6 simulations in which CED- 9/CED-4 was simulated alone and as ternary complex with EGL-1. Experimental studies show that the binding of EGL-1 to CED-9 results in the dissociation of CED-4 from CED-9 ^8, 25^. The winged-helix domain of CED-4 displays higher mobility in both simulations. However, the direction of mobility is towards the direction EGL-1 binding site and the helical domain of CED- 4a in System-5 (CED-9/CED-4) (Figure 9A). In the ternary complex in which EGL-1 is bound to CED-9 of CED-9/CED-4 complex, the direction of mobility of winged-helix domain of CED-4a is either away or in a different direction from CED-9/EGL-1 (Figure 9B). Although winged-helix domain of CED-4a does not interact with CED-9 directly, its mobility is influenced by the binding of EGL-1 to CED-9. We believe that this domain in CED-4a will play a key role in the dissociation of CED-4 from CED-9 upon binding to EGL-1. To further understand this phenomenon, we analyzed the stability of important non-covalent interactions between CED-4 and CED-9.

**Figure 8:**
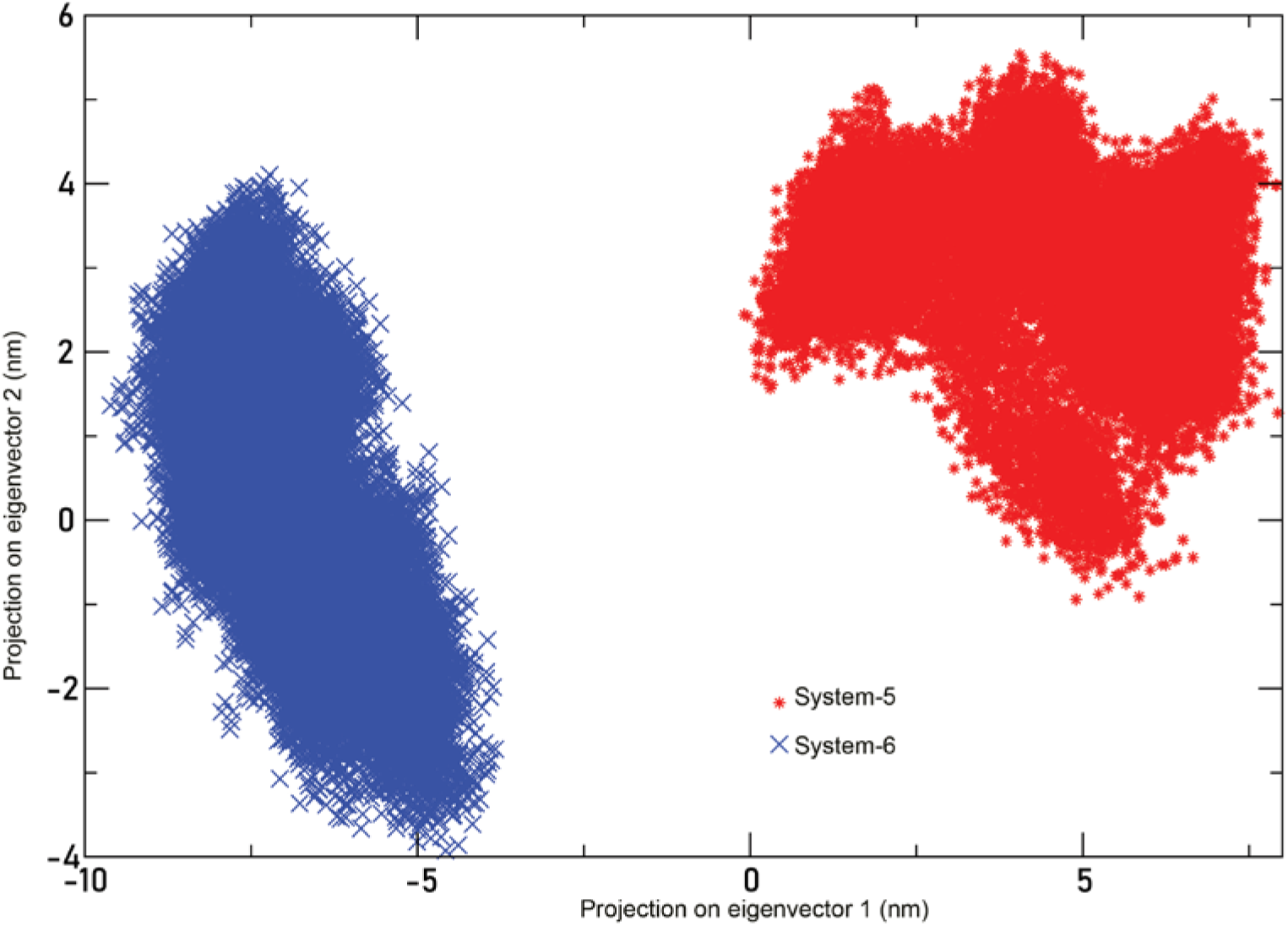
PC1 vs PC2 plot for CED-9/CED-4 complex for System 5 and System 6 simulations. This analysis was performed for the last 500 ns of production runs for both the simulations.

**Figure 9:**
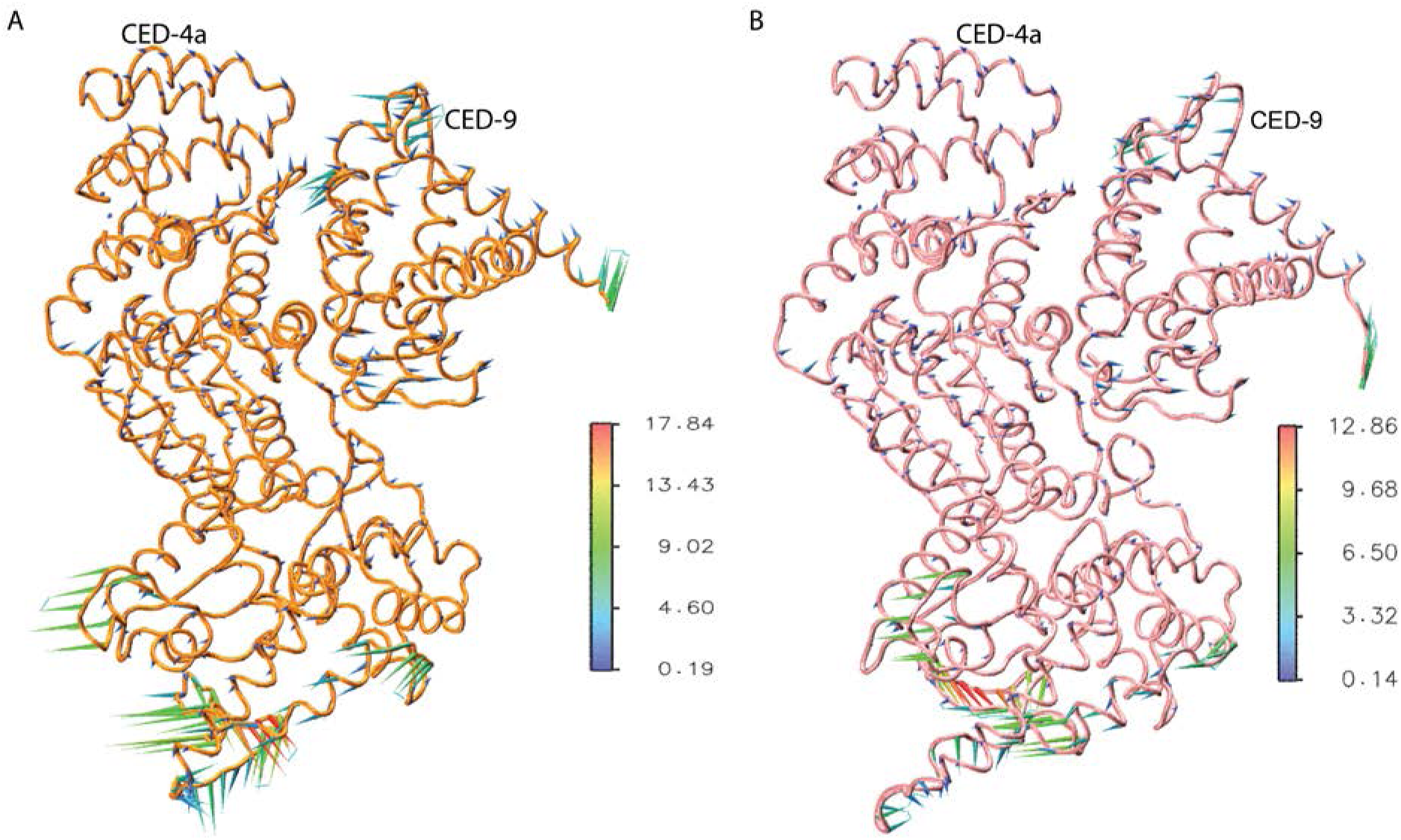
Porcupine plots of CED-9/CED-4 for the first Eigen vector (PC1) when the complex was simulated alone or with bound EGL-1.

### Non-covalent interactions between CED4 and CED9: CED-9/CED-4 vs CED-9/CED-4/EGL-1

Experimentally determined structure of CED-9/CED-4 complex has helped to identify the regions and residues in CED-9 and CED-4a that are involved in interactions between the two proteins ^25^. We determined all the residues from CED-9 and CED-4a that are within 4 Å from each other in the crystal structure. Residue range of different helical segments of CED-9 and different domains of CED-4a as defined in the PDB structures 1OHU and 2A5Y respectively are given in Tables S1 and S2. CED-9 N-terminal segment (residues 67-81), loop between helices α1 and α2 (residues 100-107), loop between α3 and α4, part of α4 (143-150) and α6 helix (residues 189-215) interact mainly with CARD and α/β domain of CED-4a. Few contacts are also found with winged helix domain of CED-4a. For the purpose of the analysis, interacting interface of CED-4a has been divided into four regions (Figure 10). Residues occurring in two helices of CARD domain (residues 22-25 and 49-65) define Region I. Both Regions II and III occur in α/β domain of CED-4a. The helix formed by residues 111-122 from α/β fold forms major part of Region II. Region III also occurs in the α/β fold and is comprised of a β-strand (186-191), an α- helix (198-212) and the subsequent loop (residues 213-223). Region IV is the small loop (367- 372) which connects helical domain and the winged helical domain of CED-4a.

**Figure 10:**
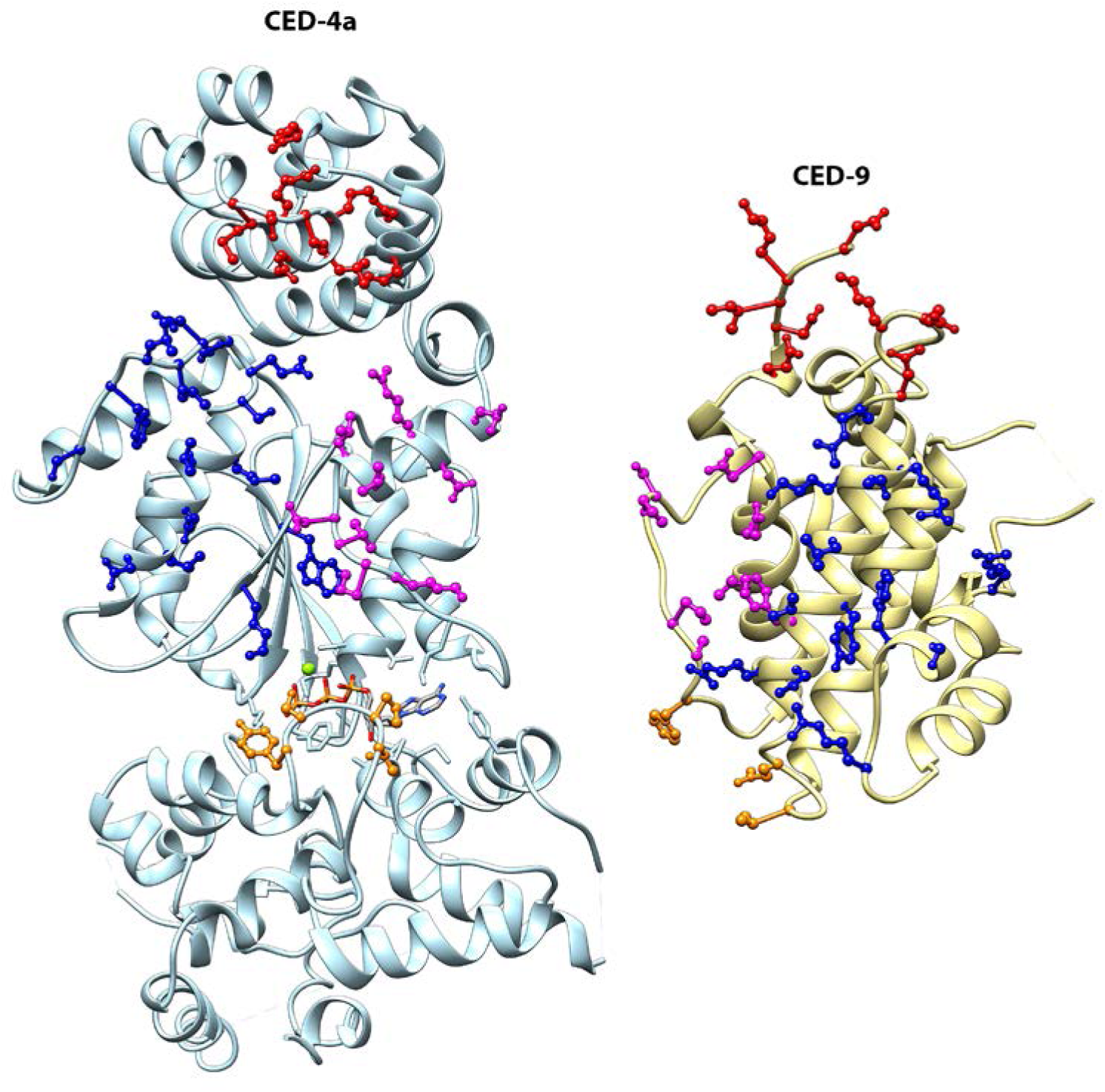
Interacting residues between CED-4a and CED-9 shown in different colors in the crystal structure (PDB ID: 2A5Y). Side-chains of interacting residues from Regions I, II, III and IV are displayed respectively in red, blue, pink and orange. A cut-off of 4 Å was used to define interacting residues between CED-4a and CED-9. For a complete list of all interacting residues in different regions, see Table S3.

We used a cut-off of 4 Å to identify residue-residue contacts between CED-9 and CED- 4a in the crystal structure and the list of residue pairs involved in interactions is provided in Table S4. The contacts were classified as salt-bridges, polar or hydrophobic contacts. Salt-bridge interactions occur between side-chain functional groups of acidic and basic residues. Polar contacts are those contacts that occur between nitrogen/oxygen of a backbone or side-chain functional group of a residue from CED-9 and nitrogen/oxygen of a backbone or side-chain functional group of a residue from CED-4a ^46^. All other contacts between CED-9 and CED-4a are considered as hydrophobic contacts. We analyzed the last 500 ns of production runs of System-5 (CED-9/CED-4) and System-6 (CED-9/CED-4/EGL-1) simulations and found out all the residue-residue contacts between CED-9 and CED-4a. The residues that are within 4.0 Å for at least 50% of the 500 ns production runs were identified. Such stable contacts helped to identify the role of some specific interactions between mammalian anti-apoptotic proteins Bcl-2 and Mcl-1 and their interacting BH3 peptides ^46, 47^. Residues in CED-9 and CED-4a that are involved in contact within 4 Å for at least 50% of the last 500 ns production run are summarized for both System-5 (CED-9/CED-4) and System-6 (CED-9/CED-4/EGL-1) simulations (Table 3) and are shown in Figure 11. We have analyzed the residue-residue contacts in the crystal structure and compared them with the stable contacts found in the two simulations. This will indicate whether the binding of EGL-1 has weakened the interactions between CED-9 and CED- 4a and to what extent the EGL-1 binding has affected the binding of CED-9 with CED-4a. Our analysis is presented for each of the four regions. The comparative analysis was carried out for CED-9/CED-4 complex with (System-6) and without EGL-1 (System-5). Our PCA analysis shows that EGL-1’s association with CED-9/CED-4 makes the complex to access distinct conformations. Although winged-helix domain of CED-4a is not directly involved in interactions with CED-9, the porcupine plots reveal large motions in winged-helix domain of CED-4a. However, the direction of motion is different when EGL-1 binds to CED-9/CED-9 complex. We have examined to find whether there exists a correlation between the winged-helix motion of CED-4a and the stability of non-covalent interactions between CED-9 and CED-4a.

**Figure 11:**
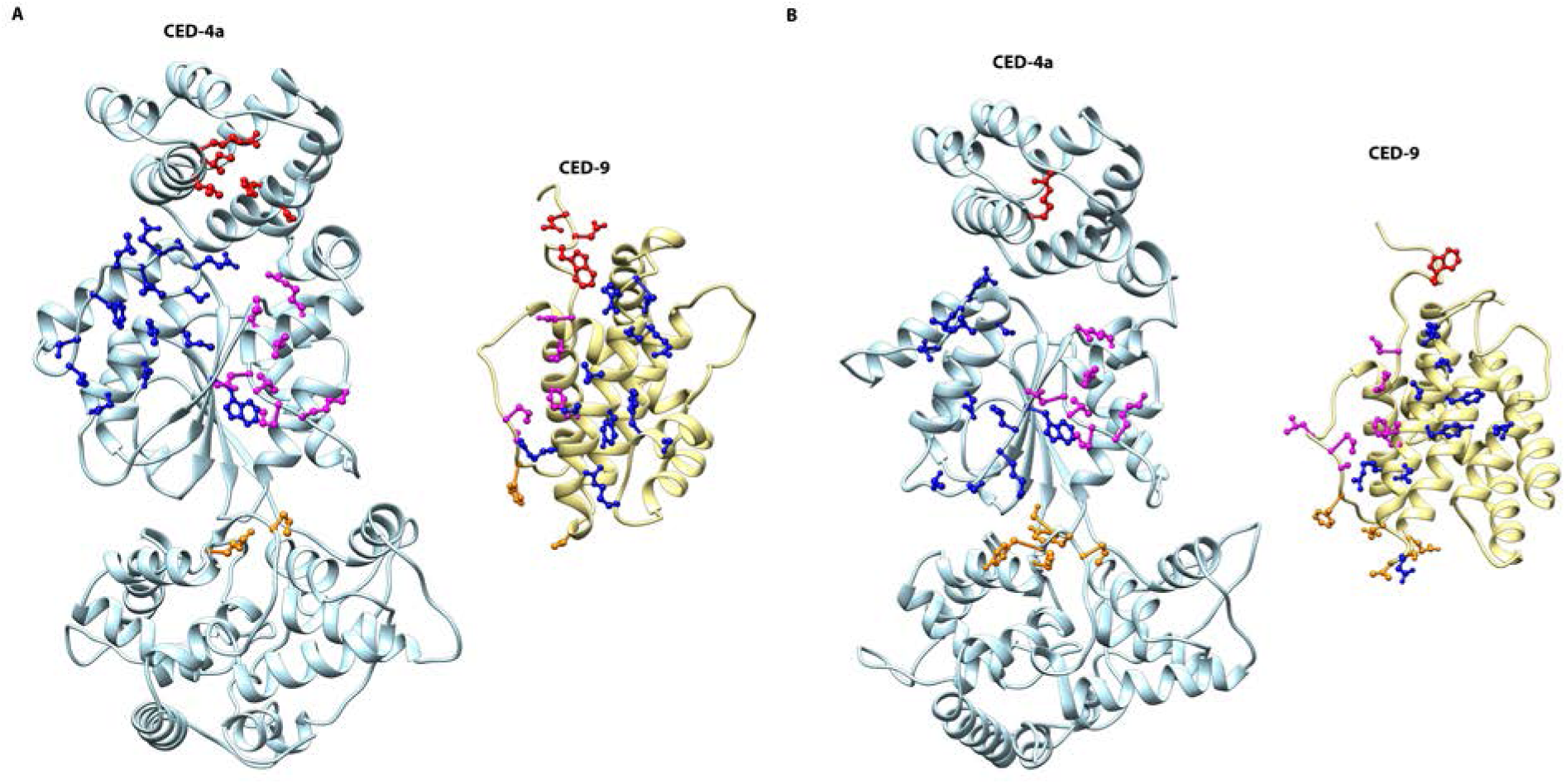
Interacting residues involved in stable contacts from different regions of CED-4a and the corresponding interacting residues from CED-9 are shown for (A) System-5 (CED-9/CED-4) and (B) System-6 (CED-9/CED-4/EGL-1) simulations.

**Table 3:** Residue pairs involved in stable non-covalent interactions between CED-9 and CED-4a in System-5 (CED-9/CED-4) and System-6 (CED-9/CED-4/EGL-1) simulations

| CED-4a Region<sup>a</sup> | Non-covalent interaction<sup>b</sup> | System-5 (CED-9/CED-4)<sup>c</sup> | System-6 (CED-9/CED-4/EGL-1)<sup>b</sup> |
| --- | --- | --- | --- |
| Region-I | Salt-bridge | D72...R50 | No stable salt-bridge |
|  | Polar | N71...T15, N71...H14, D72...L51 | No stable polar contacts |
|  | Hydrophobic | W73...L51, W73...H14, W73...I54 | W73...R50 |
| Region-II | Salt-bridge | D79...R117 | D79...R117, E81...R117 |
|  | Polar | E81...L120, E81...L121, A102...K126, R196...N123, G104...K126 | P103...N123, G104...K126 |
|  | Hydrophobic | G82...L120, G101...P125, A102...P125, P103...L120, P103...V124, P103...N123, R196...P125, F199...N123, F199...L120 | G101...P125, A102...P125, P103...L120, P103...V124, G104...V124, G104...P125, L105...K126, R196...N123, F199...N123, V200...N123 |
| Region-III | Salt-bridge | R143...D206, R211...D216 | R211...D216 |
|  | Polar | R143...D202, S147...V223, S147...S222, L204...S213, T208...D215, R209...L218, R211...E214, R211...D215, N212...D216 | V189...S193, E190...S193, R196...K191, N197...D206, L204...S213 |
|  | Hydrophobic | F146...L209, F146...F220, R196...W189, V200...M210, Y201...L209, Y201...D206, L204...L217, L204...L209, T208...D216, N212...L218 | R154...N219, E190...T195, G193...S193, R196...L190, R196...W189, Y201...L218, L204...L217, L204...L218, F205...L218, T208...D216 |
| Region-IV | Salt-bridge | No stable salt-bridge | No stable salt-bridge |
|  | Polar | No stable polar contacts | G96...K372, F100...Y369, F100...S370, E187...Y371 |
|  | Hydrophobic | F100...P368, V789...K372 | F100...P368, S188...Y371, V189...Y371, Q192...S370, Q192...Y371 |

<sup>a</sup>For residues that constitute different regions of CED-4a, see Table S3
<sup>b</sup>For definition of different non-covalent interactions, see the main text.
<sup>c</sup>For each interacting residue pair, the first and second residues correspond to CED-9 and CED-4a respectively. Residue numbering for CED-9 and CED-4a is according to the PDB IDs 1OHU and 2A5Y respectively.

<u>Region I</u>: In the crystal structure, Asp-67 in the N-terminus of CED-9 is simultaneously involved in salt-bridge interactions with Arg-24 and Arg-53 of CARD domain of CED-4. Residues from the N-terminal regions (residues 67 to 71 and 74) and the C-terminal end of α6 helix (residues 211, 212, 215 and 216) of CED-9 make polar and hydrophobic contacts with the CARD domain helix formed by residues 49-65. In total 9 polar and 11 hydrophobic contacts are found in the crystal structure (Table S4). However in System-5 (CED-9/CED-4) simulation, we found only 3 polar and 3 hydrophobic contacts that can be described as stable contacts (Table 3). The salt-bridges formed by Asp-67 of CED-9 are no longer stable in System-5 (CED-9/CED-4) simulation. Instead, a new salt-bridge is formed between Asp-72 of CED-9 and Arg-50 of CED-4. In the case of System-6 (CED-9/CED-4/EGL-1) simulation, majority of these interactions are destabilized (Table 3). We have found just one stable contact between Trp-73 of CED-9 and Arg-50 of CED-4 and this can be described as hydrophobic contact. Compared to the crystal structure, interactions between CED-9 and the residues involving Region-I of CED-4a have weakened in CED-9/CED-4 complex to some extent. However, almost all interactions involving residues from Region-I of CED-4 are mostly destabilized in System-6 (CED-9/CED-4/EGL-1).

<u>Region II</u>: There is one salt-bridge found between the residues Asp-79 of CED-9 and Arg-117 of CED-4a in the crystal structure. This region of CED-4a has large number of hydrophobic contacts with CED-9. There are 20 hydrophobic contacts and only 5 polar contacts in the crystal structure (Table S4). In System-5 (CED-9/CED-4) simulation, the salt-bridge between Asp-79 of CED-9 and Arg-117 of CED-4 is maintained. However, the number of stable hydrophobic contacts has dropped to 9 from the 20 contacts observed in the crystal structure (Table 3). The number of stable polar contacts remains same although these contacts now occur between different pairs of residues. In System-6 (CED-9/CED-4/EGL-1) simulation, in addition to the salt-bridge observed between Asp-79 of CED-9 and Arg-117 of CED-4a, an additional new salt-bridge is formed between Glu-81 of CED-9 and Arg-117 of CED-4 and this interaction is found to be stable. However, the number of stable polar contacts has come down to 2 and the stable hydrophobic contacts are found in only 10 residue pairs (Table 3). Thus in both System-5 (CED-9/CED-4) and System-6 (CED-9/CED-4/EGL-1) simulations, the number of stable hydrophobic contacts has come down to about 50% of those found in the crystal structure involving residues from Region-II of CED-4a. CED-9/CED-4/EGL-1 complex has resulted in less number of stable polar contacts compared to those found in System-5 (CED-9/CED-4) simulation indicating again that the interaction of EGL-1 with CED-9 has further weakened the stable interactions between CED-9 and CED-4a.

<u>Region-III</u>: In the CED-9/CED-4 crystal structure, two salt-bridges, 9 polar contacts and 16 hydrophobic contacts have been identified (Table S4). Arg-143 and Lys-207 of CED-9 participate in salt-bridge interactions respectively with Asp-206 and Glu-214 of CED-4a.

However, only the salt-bridge between Arg-143 and Asp-206 is found to be stable in System-5 (CED-9/CED-4) simulation (Table 3). An additional stable salt-bridge is formed and is determined to be stable between Arg-211 of CED-9 and Asp-216 of CED-4a in System-5 (CED9/CED-4) simulation. The number of stable polar contacts in System-5 (CED-4/CED-9) is the same as found in the crystal structure. However although the number is same, the residue pairs participating in polar contacts are different. The number of stable hydrophobic contacts has come down from 16 in the crystal structure to 11 in System-5 (CED-9/CED-4) simulation. The salt-bridge between Arg-211 of CED-9 and Asp-216 of CED-4a is found to be stable in System- 6 (CED-9/CED-4/EGL-1) simulation also (Table 3). However, the stable polar contacts are found only in five residue pairs. Similarly only 10 stable hydrophobic contacts are observed in CED-9/CED-4/EGL-1 simulation indicating that EGL-1’s interaction with CED-9 has diminished the interaction strength of CED-9 and CED-4a.

<u>Region IV</u>: Only a small number of residues are involved in interaction between residues of Region-IV in CED-4a and CED-9. This includes 1 polar contact and 4 hydrophobic contacts in the crystal structure (Table S4). In System-5 (CED-9/CED-4) simulation, no polar contacts are found to be stable. Only two hydrophobic contacts are stable (Table 3). This is the only region which has resulted in increased number of stable contacts in System-6 (CED-9/CED-4/EGL-1) simulation. We have found 4 polar contacts and 5 hydrophobic contacts that remain stable during the last 500 ns of production run in CED-9/CED-4/EGL-1 simulation (Table 3).

In summary when we consider all the four regions of CED-4 together, the binding of EGL-1 to CED-9 has reduced the number of salt-bridge and polar interactions between CED-9 and CED-4a. Only three stable salt-bridges are found in System-6 (CED-9/CED-4/EGL-1) compared to 4 in System-5 (CED-9/CED-4) simulation and 5 in the crystal structure. The number of stable polar contacts has drastically come down to 11 in System-6 (CED-9/CED- 4/EGL-1) simulation compared to 17 in System-5 (CED-9/CED-4) and 24 in the crystal structure. Only half of hydrophobic contacts found in the crystal structure are found to be stable in both System-5 and System-6 simulations. Analysis of stable non-covalent interactions between CED-9 and CED-4a clearly reveal that EGL-1 has significant impact upon binding to CED-9 to reduce CED-9’s affinity to CED-4a. Binding of EGL-1 has compromised a large number of polar and hydrophobic interactions that occur between CED-9 and CED-4a.

## Discussion

Binding of EGL-1 to CED-9/CED-4 complex and subsequent dissociation of CED-4 from CED-9 is the most important step in the linear pathway of programmed cell death in nematodes. The molecular mechanism of how EGL-1 after binding to CED-9 releases CED-4a from the CED-9/CED-4 complex is not known. While the previous studies have succeeded in determining the crystal structures of CED-9 in apo ^8^ or in complex with CED-4 ^25^ or EGL-1 ^9^, the ternary complex structure of CED-9/CED-4/EGL-1 is yet to be determined experimentally. This may be due to the transient nature of the ternary complex. The primary aim of this study is to understand the molecular events that lead to the dissociation of CED-4 from CED-9 upon EGL-1’s binding. To achieve this goal, we first constructed a model of the ternary complex involving all three molecular players. The experimentally determined structures of CED-9 in complex with EGL-1 and CED-4 were used to build CED-9/CED-4/EGL-1 ternary complex. We considered six different systems all involving CED-9 either in the apo form or in complex with EGL-1 and/or CED-4 (Table 1). We performed molecular dynamics simulations of all six systems for a period of 500 ns to 1 μs to understand the dynamics of CED-9 and CED-4 in different conditions. CED-9 is present in all the systems studied. Both 1D- and 2D-RMSD analyses demonstrated that CED-9 displays conformational heterogeneity and its dynamics is influenced by its interaction with EGL-1 and/or CED-4. While CED-4’s association with CED-9 is through its asymmetric dimer (CED-4a and CED-4b), comparative MD simulations of CED-4 complex structures with and without CED-4b illustrate the role of CED-4b in regulating the flexibility of CED-4a. CED-4a has two interacting interfaces, one with EGL-1 and another with CED-4b. The α/β fold and the helical fold of CED-4b forms the interface with the CARD, α/β domain and winged-helix domain of CED-4a. When CED-4a is simulated with CED-9 without CED-4b, CED-4a exhibits higher conformational flexibility due to the absence of many interactions with CED-4b (Figure 5). The plot of PC1 versus PC2 indicates distinct conformational changes when CED-9 binds to EGL-1 and/or CED-4 (Figure 7). Similarly the CED-9/CED-4 complex as a whole displays distinct dynamics when it is bound to EGL-1 as revealed by the PC1 versus PC2 plot (Figure 8).

To further make sense of these conformational changes that will eventually disrupt CED- 9’s interaction with CED-4a, we analyzed the non-covalent interactions between CED-4a and CED-9 found in the crystal structure. Similarly, we determined stable non-covalent contacts that occur between CED-9 and CED-4a with and without EGL-1. Our analysis shows that the number of salt-bridges, polar and hydrophobic contacts has significantly weakened upon EGL-1 binding in comparison to CED-9/CED-4 complex without EGL-1 (Table 3). This clearly indicates that binding of EGL-1 has consequences for the CED-9 binding to CED-4a and will eventually disrupt the CED-9’s association with CED-4a. This study also revealed the likely steps that will occur during the dissociation of CED-4a. Among the four regions of CED-4a that form the interface between the CED-4a and CED-9, interactions involving the residues of Region-I are the first to be destabilized. Salt-bridge and polar interactions due to Region-II and Region-III residues are mostly weakened in CED-9/CED-4/EGL-1 ternary complex. Surprisingly, new stable interactions occur between the loop connecting the helical and winged-helix domains of CED-4a and CED-9 residues. It is tempting to speculate that these interactions stabilize an intermediate conformational state of CED-9/CED-4 complex before CED-4a completely breaks away from CED-9. Our study first time has given rise to a hypothesis that the winged-domain of CED-4a is likely to be involved in the disruption of interactions between CED-9 and CED-4a. Experimentally substituting some of these residues identified in this study in winged-helix region will confirm the role of winged helix domain in modulating the CED-9/CED-4 complex structure.

Although we have performed long simulations for a period of at least 1 μs to understand the dynamics of CED-4/CED-9 interactions with and without bound EGL-1, our simulations have not resulted in complete dissociation of CED-4 from CED-9 in the ternary complex. Our studies, however, have clearly shown that the interactions between CED-4a and CED-9 have greatly weakened as a result of EGL-1’s binding to CED-9. There could be several reasons for this observation. It is quite possible that even longer simulation is required to see a complete disruption of interactions between CED-9 and CED-4a. It is also possible that some of the non-covalent interactions are not correctly represented in the currently available force-fields. There are reports that the force-fields such as AMBER and CHARMM ^48, 49^ might have overestimated the attractive interactions due to non-covalent contacts ^50^. In protein folding studies, it has been shown that the strength of salt-bridges and hydrophobic interactions have been stronger ^51–54^ resulting in protein aggregates. Nevertheless, the current study has unambiguously demonstrated that the binding of EGL-1 loosens the interactions between CED-9 and CED-4a that will eventually result in the release of CED-4a. We have also found the possible involvement of winged-helix domain in CED-4a which is not in direct contact with EGL-1. Results of the present study can be used to design new mutation experiments in winged-helix domain of CED- 4a to understand the cross talk between the three important molecular players of nematode apoptosis CED-9, CED-4 and EGL-1.

## Supporting information

Supplementary Materials with Tables and Figures

CED-9/CED-4/EGL-1 ternary complex model in PDB format

## Acknowledgements

We gratefully acknowledge the High Performance Computing Facility at IIT-Kanpur. RS was Pradeep Sindhu Chair Professor during the course of this work. CNR thanks BINC Fellowship. We thank our lab members for useful discussion.

## Supplementary Material

Tables defining the different helical region of CED-9, different domains of CED-4, regions of CED-4 interacting with CED-9 and non-covalent interactions between CED-9 and CED4 found in the crystal structure of CED-9/CED-4 complex are provided. RMSF plot of CED-9 with the CED-9 reference structure taken from CED-9/EGL-1 complex and 2D-RMSD plots of System-3 and System-4 are available. The coordinates of the modeled CED-9/CED- 4/EGL-1 ternary complex in PDB format can be obtained from the Supporting Information.

## Notes

### Competing Interest Statement

The authors have declared no competing interest.

