## Supplementary Materials with Tables and Figures for "Dissociation of CED-4 from CED-9 upon EGL-1 binding: Molecular mechanism of linear apoptotic pathway in *Caenorhabditis elegans*"

**Table S1**: Major helical regions in CED-9 as defined in the CED-9 apo crystal structure (PDB ID: 1OHU)

| S. No. | Helix | Residue range |
| --- | --- | --- |
| 1. | α1 | 79-94 |
| 2. | α2 | 110-127 |
| 3. | α3 | 127-139 |
| 3. | α4 | 145-154 |
| 4. | α5 | 167-187 |
| 5. | α6 | 194-218 |
| 6. | α7 | 220-241 |

**Table S2**: Different domains of CED-4 as defined in the crystal structure of CED-9/CED-4 complex (PDB ID: 2A5Y)

| Domains in CED-4 | Residue range |
| --- | --- |
| CARD domain | 1-105 |
| Three-layered α/β fold | 106-290 |
| Helical domain | 291-370 |
| Extended winged helix domain | 371-549 |

**Table S3**: Residues of CED-4 from the four different regions that are involved in interactions with CED-9

| CED-4 interacting Region | Interacting residues of CED-4 with CED-9^a^ | CED-9 residues interacting with residues from one of the CED-4 regions^b, c^ |
| --- | --- | --- |
| Region-I | 14, 15, 18, 24, 47 to 54 | 67 to 71, 74, 211, 212, 215, 216 |
| Region-II | 113, 116, 117, 120, 121, 123 to 126 | 79, 81, 82, 85, 101 to 105, 106, 107, 196, 199, 200 |
| Region-III | 154, 189, 190, 191, 193, 195, 202, 203, 206, 209, 210, 213 to 220, 222, 223 | 143, 146, 147, 154, 189, 190, 193, 196, 197, 200, 201, 204, 205, 207 to 209, 211, 212 |
| Region-IV | 367 to 372 | 96, 100, 187 to 189, 192 |

^a^Residues are listed as interacting residues if they interact with CED-9 residues either in the crystal structure or if they are involved in stable contacts in System-5 (CED-9/CED-4) or System-6 (CED-9/CED-4/EGL-1) simulations. For details of stable contact, see the main text.

^b^Residues are listed as interacting residues if they interact with CED-4 residues from one of the four regions either in the crystal structure or if they are involved in stable contacts in System-5 (CED-9/CED-4) or System-6 (CED-9/CED-4/EGL-1) simulations. For details of stable contact, see the main text.

^c^Residues 189, 196 and 200 from CED-9 are involved in interactions with residues from more than one region of CED-4

**Table S4**: Non-covalent interactions between CED-9 and CED-4 in the crystal structure

| CED-4a Region^a^ | Non-covalent interactions^b^ | CED-9-CED-4a Interacting residue pairs^c^ |
| --- | --- | --- |
| Region-I | Salt-bridge | D67…R24, D67…R53 |
|  | Polar | D67…S48, G68…S48, G68…R53, N71…T49, E74…T49, E74…E52, R211…E52, N212…S48, N212…E52 |
|  | Hydrophobic | G68…R50, K69…R50, I70…I18, I70…R50, I70…L51, E74…L51, R211…M47, N212…M47, K215…T49, E216…S48 |
| Region-II | Salt-bridge | D79…R117 |
|  | Polar | A102…K126, G104…V124, P106…Q113, S107…Q113, R196…N123 |
|  | Hydrophobic | D79…L121, E81…L121, E81…L120, G82…L120, V85…L120, G101…P125, A102…P125, P103…L120, P103…N123, P103…V124, P103…P125, G104…L120, G104…K126, P106…D116, P106…R117, P106…L120, R196…P125, F199…L120, F199…N123, V200…N123 |
| Region-III | Salt-bridge | R143…D206, K207…E214 |
|  | Polar | R143…D202, R154…L218, R196…W189, N197…D206, Y201…D206, K207…S213, R211…S213, R211…E214, R211…D215 |
|  | Hydrophobic | R143…L203, F146…L209, F146…F220, S147…S222, G193…K191, V200…L209, V200…M210, Y201…L209, L204…L209, L204…S213, L204…L217, T208…D215, T208…D216, T208…L217, T208…L218, R209…L218 |
| Region IV | Salt-bridge | No salt-bridge found |
|  | Polar | Q192…S370 |
|  | Hydrophobic | F100…T367, F100…P368, V189…S370, V189…Y371 |

^a^For different regions of CED-4a, see Table S3

^b^For definition of different non-covalent interactions, see the main text.

^c^For each interacting residue pair, the first and second residues correspond to CED-9 and CED-4a respectively. Residue numbering for CED-9 and CED-4a is according to the PDB IDs 1OHU and 2A5Y respectively

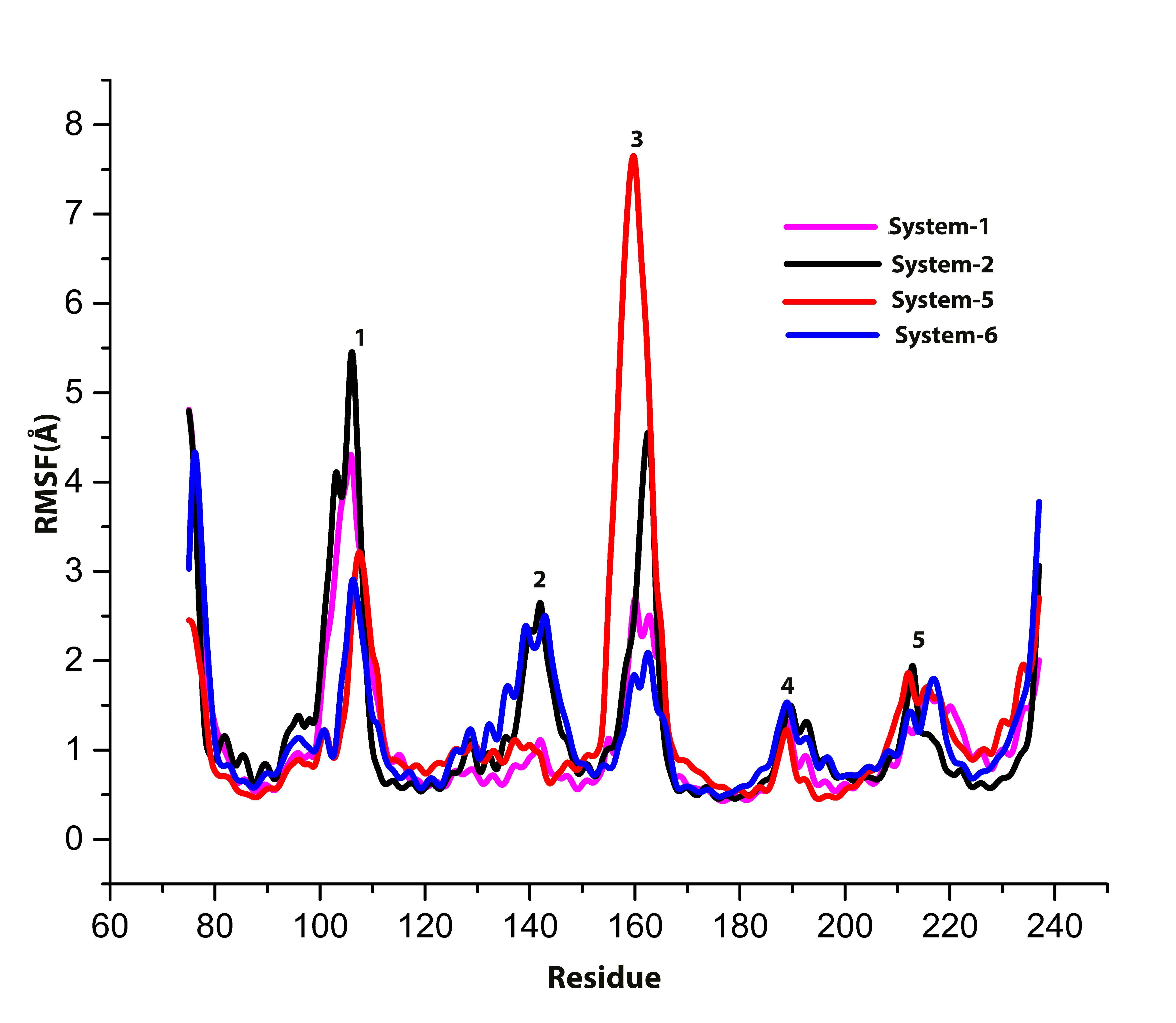

**Figure S1**: Root mean square fluctuation (RMSF) of CED-9 in different simulated systems calculated with CED-9/EGL-1 as the reference structure. The CED-9 regions that undergo large fluctuations are the loops connecting helices (1) α1 and α2 (95-109), (2) α3 and α4 (residues 130-150), (3) α4 and α5 (residues 155-166), (4) α5 and α6 (residues 185-194) and (5) α6 and α7 (residues 205-220). The pattern of fluctuations is same as that observed in Figure 3.

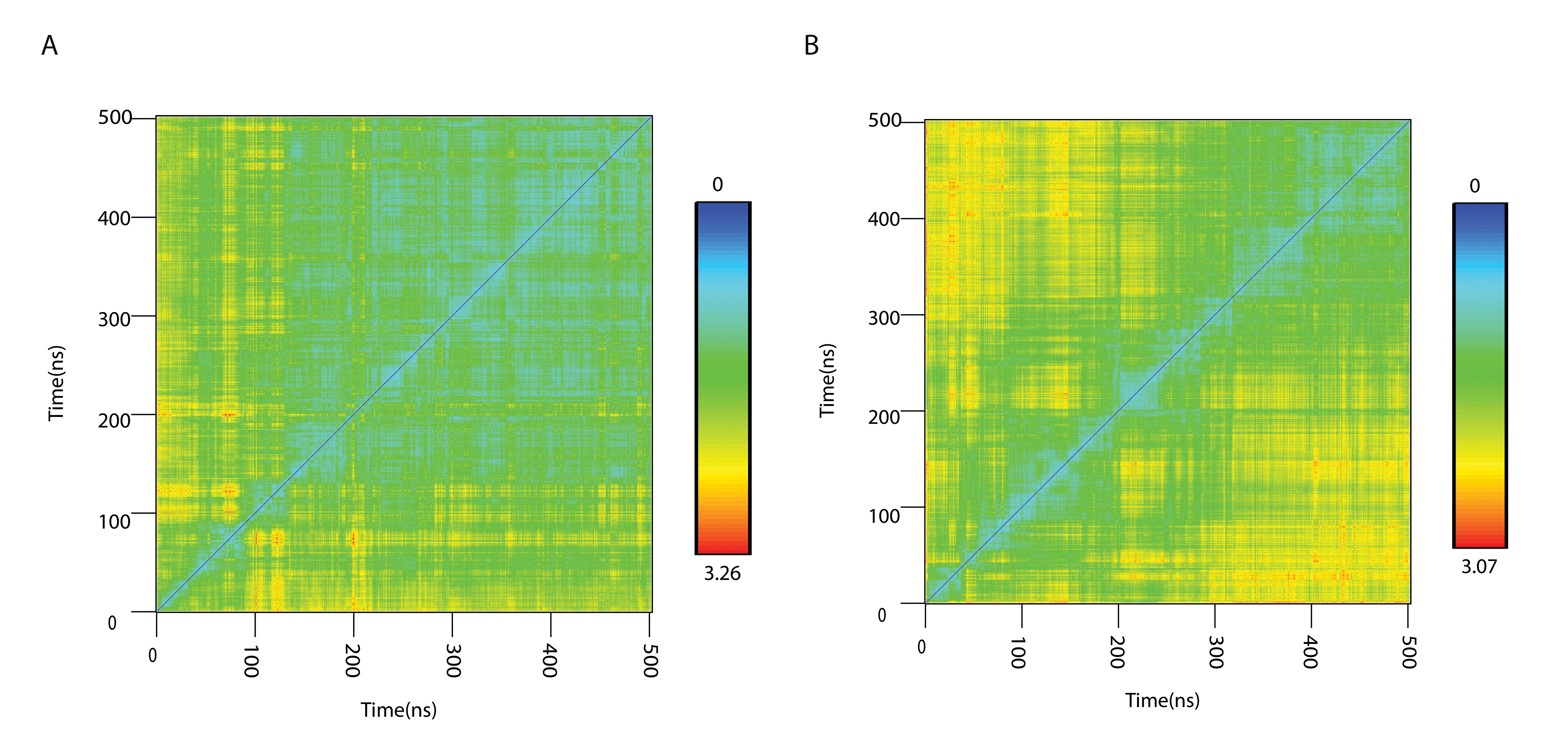

**Figure S2**: 2D-RMSD plots of CED-9 shown for (A) System-3 (CED-9/CED-4a) and (B) System-4 (CED-9/CED4a/EGL-1) simulations as a function of time. RMSD values (in Å) are displayed from blue to red covering the range of lowest to highest values.
